# Temporal Control of Decidual Inflammation by HOXA10 is Essential for Implantation and its Dysregulation is Associated with Early Pregnancy Loss

**DOI:** 10.64898/2025.12.02.691844

**Authors:** R Sharma, B Negi, R Ponsankaran, S Patil, G Godbole, A Mishra, S Shyamal, D Modi

## Abstract

Successful implantation requires precisely timed endometrial inflammation. Although an initial inflammatory burst is essential for implantation, this response must be rapidly resolved for placentation and pregnancy to progress. The mechanisms that coordinate this temporal switch remain poorly understood. Here, we identify the transcription factor HOXA10 as a key regulator of inflammatory transitions in decidual stromal cells and examine how its dysregulation contributes to implantation failure and early pregnancy loss.

In mice, HOXA10 expression decreases transiently at implantation and rises again post-implantation. Silencing HOXA10 in human decidualized stromal cells induced a robust pro-inflammatory state, altered integrin and cytoskeletal gene expression, and impaired stromal substrate adhesion. In non-pregnant Hoxa10 hypomorphs, stromal cells exhibited elevated IL1&#946; and TXNIP, activation of the NLRP3/ASC inflammasome, and dysregulation of receptivity markers. During pregnancy, persistent HOXA10 deficiency prevented the resolution of inflammation, resulting in disorganized decidua, defective placentation, infertility, or progressive reproductive decline.

To assess translational relevance, we analyzed publicly available bulk and single-cell RNA-seq datasets from first-trimester human decidua and from women with recurrent pregnancy loss (RPL). Single-cell analysis revealed that HOXA10 is selectively low in inflammatory decidual stromal cell clusters, which display concordant upregulation of IL1B, PYCARD, TXNIP, and inflammasome components. Consistently, both bulk and single-cell datasets from women with RPL showed reduced HOXA10 accompanied by increased inflammatory gene expression.

Overall, HOXA10 functions as a temporal switch that enables the initial inflammatory activation required for implantation and subsequently suppresses inflammation to support decidual organization, adhesion, and placentation. Loss of this switch leads to persistent decidual inflammation and is associated with pregnancy loss in both mouse models and humans.

**Graphical abstract:** 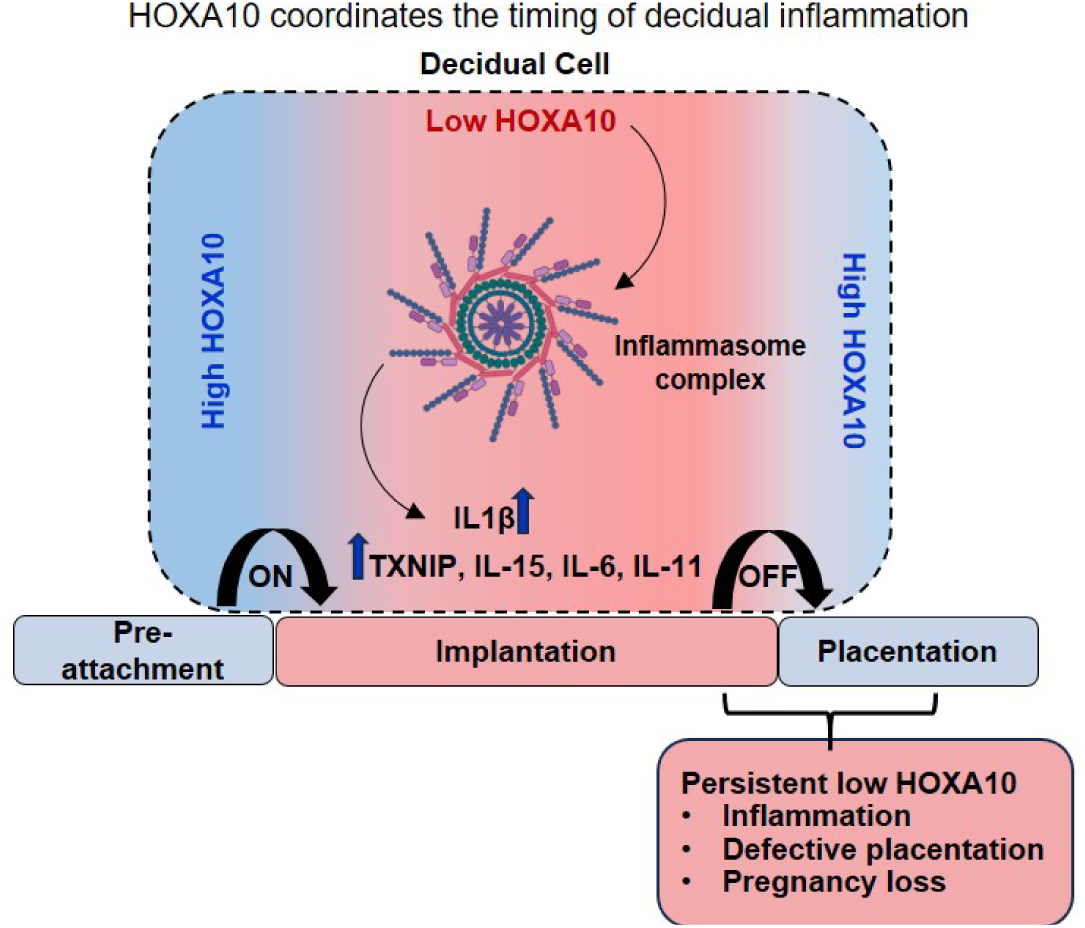

## Introduction

The endometrium is the inner lining of the uterus where embryo implantation occurs. Under most circumstances, it remains refractory to implantation except during a brief, hormonally regulated window of receptivity. Once considered a passive scaffold for the implanting embryo, the endometrium is now recognized as a dynamic tissue that actively assesses embryonic quality and determines whether to support or terminate implantation [1,2]. This bidirectional embryo–endometrial communication underlies the “selector” function of the endometrium, allowing it to distinguish between high- and poor-quality embryos and modulate implantation responses accordingly [1,3].

A key feature of the receptive endometrium is the transformation of stromal fibroblasts into decidual cells. This decidualization process is essential for embryo implantation, placental development, and the maintenance of pregnancy. Impaired decidualization contributes to infertility, recurrent implantation failure (RIF), recurrent pregnancy loss (RPL), and other obstetric complications [4,5]. Recent single-cell RNA sequencing (scRNA-seq) studies have revealed that the decidua contains multiple stromal cell subtypes [6,7], including glycolytic, myofibroblastic, and an inflammatory subset known as inflammatory decidual stromal cells (iDSCs). Although the exact functions of these subpopulations are still emerging, iDSCs are thought to contribute to the inflammatory environment required for successful implantation. Although pregnancy is broadly viewed as an anti-inflammatory state, implantation paradoxically begins with a tightly regulated pro-inflammatory burst [8–10]. This transient inflammatory response facilitates trophoblast invasion and vascular remodeling but must be rapidly resolved to sustain pregnancy [11–13]. Comparative studies in eutherian mammals suggest that the evolution of decidualization enabled precise control of this inflammatory window, supporting successful implantation and placentation [9,13–15]. Dysregulation of this inflammatory balance by either excessive or insufficient is implicated in RPL, RIF, endometrial disorders, and other pregnancy complications [1,3,4,6,16–18].

Despite its importance, the molecular mechanisms that regulate the inflammatory phenotype of decidual stromal cells remain poorly understood. In particular, it is unclear what drives the emergence of inflammatory decidual subsets and how this process is temporally coordinated around implantation. HOXA10, an Abdominal-B class homeobox transcription factor, is a central regulator of uterine development and function [19–21]. In adult tissues, HOXA10 is highly expressed in the endometrium and is essential for establishing receptivity and progesterone-driven decidualization [19]. Loss of HOXA10 impairs stromal proliferation, disrupts decidual transformation, and has been linked to multiple reproductive pathologies [19,22–24]. Intriguingly, while HOXA10 is critical for establishing receptivity, its expression declines in the decidua after implantation in both human and non-human primates [25,26]. Furthermore, silencing HOXA10 in decidualized human stromal cells induces pro-inflammatory cytokines such as IL11, IL15, LIF, and IL6 [26,27].

These findings led us to hypothesize that HOXA10 not only supports decidualization but also regulates the inflammatory programs of decidual stromal cells. Specifically, we hypothesized that a transient reduction in HOXA10 during implantation may allow the emergence of inflammatory decidual cells, while its re-expression after implantation is required to resolve inflammation and sustain pregnancy. In this study, we investigate the role of HOXA10 in controlling endometrial inflammation across human and murine models and explore its relevance in reproductive failure, including RPL.

### Materials and Methods

#### Human and Animal Ethics

All animal procedures were approved by the Institutional Animal Ethics Committee (IAEC) of ICMR–NIRRCH (project numbers 08/18 and 07/21). Collection of human endometrial tissues for primary stromal culture was approved by the Institutional Ethics Committee for Human Studies, using the same donor samples described previously [26–28].

#### Development of *HOXA10* hypomorphs and fertility assessment

*HOXA10* hypomorphic mice were generated by testicular transgenesis using a shRNA construct, as described earlier [22]. Animals were genotyped, bred, and maintained under standard environmental conditions (12-h light–dark cycle) in the ICMR–NIRRCH animal facility. Regularly cycling, three-month-old C57BL/6 wild-type females and Hoxa10 hypomorphs were housed with proven fertile wild-type males. The day a vaginal plug was observed was designated as Day 1 of pregnancy.

Uteri from wild-type and hypomorphic mice were collected either at diestrus or at specified gestational time points. For fertility assessment, mating outcomes including date of delivery and number of pups born per mating cycle were recorded prospectively.

#### *In vitro* decidualization and *HOXA10* silencing

Human endometrial stromal cells were obtained as described previously [26–28]. Briefly, proliferative phase endometrium was collected from women after informed consent. They were aged 30–46 years with regular menstrual cycles (28–35 days) who had not used steroid hormones for at least three months. Stromal cells were isolated, assessed for purity, and validated [27]. Cells from two donors were pooled to generate three biological replicates.

Decidualization was induced by treating stromal cells with 17β-estradiol (10[[M) and progesterone (10[[M) for 20 days [27]. Decidualization efficiency was confirmed by measuring the mRNA expression of *PRL* and *IGFBP1* in the cells [28].

For *HOXA10* silencing, decidualized stromal cells (Day 20) were transfected with *HOXA10*-specific siRNA (Supplementary Table 1) or scrambled siRNA (control) using HiPerFect reagent (both from Qiagen). Total RNA from scrambled and *HOXA10* siRNA-transfected cells was extracted using TRIzol (Thermo Fisher Scientific) at 24, 48, and 72 hours post-transfection. *HOXA10* mRNA levels were quantified at 24, 48, and 72 hours post-transfection by real-time PCR [27].

#### Microarray and Data Analysis

cDNA was labelled with Cy3 or Cy5 dyes using the Indirect Labeling and Cleanup System (Promega). Labelled samples were co-hybridized to a 33K Human Oligo microarray. Each time point included triplicate biological samples and one dye-swap, yielding four arrays per time point. Microarray data were globally normalized, and genes with log[fold change >2 and p < 0.05 were considered differentially expressed. GO and pathway analyses were performed using WebGestalt (https://www.webgestalt.org/), and results were visualized in RStudio, (http://www.rstudio.com/), SR plots (https://www.bioinformatics.com.cn/en), and Cytoscape (https://cytoscape.org/). The data is available with the accession ID GSE35801.

#### Real-time PCR

Real-time PCR was performed with SYBR green super mix (Bio-Rad Laboratories) on the CFX96 Real-Time PCR system (Bio-Rad) following the protocol described previously [27]. Gene expression levels were normalized to *18s rRNA*. Primer sequences, annealing temperatures, and product sizes are listed in the Supplementary Table 1.

#### Cytokine profiling by multiplex immunoassay

Culture supernatants from decidualized and *HOXA10*-silenced stromal cells were collected at 48 h and 72 h as described earlier [26]. Cytokine levels were measured using the Bio-Plex Pro Human Cytokine Assay (Bio-Rad) according to manufacturer instructions. Data were expressed as mean concentrations or as fold change relative to scrambled siRNA controls.

#### Adhesion assays

Control and *HOXA10* knockdown cells were seeded onto extracellular matrix–coated plates (gelatin, fibronectin, laminin, collagen, poly-L-lysine from Merck and Matrigel from BD Biosciences). After incubation, non-adherent cells were removed by gentle washing. Adherent cells were quantified colorimetrically using the Crystal Violet method [29], and absorbance values were normalized to control cells.

#### Immunofluorescence/ Immunohistochemistry

Paraffin-embedded uterine tissues were sectioned at 5 µm, deparaffinized, rehydrated, and either stained with hematoxylin and eosin (H&E) or processed for immunostaining. H&E sections were mounted in DPX and imaged by bright-field microscopy (Olympus).

Immunofluorescence or immunohistochemistry was performed as detailed previously [22,30,31]. Briefly, the sections underwent antigen retrieval and blocking. Immunofluorescence for HOXA10, IL1β, TXNIP, NLRP3, and ASC was performed using Tyramide Signal Amplification (PerkinElmer or Biotium). After each TSA reaction, slides were re-retrieved and incubated with the next antibody. Sections were mounted with EverBrite™ Hardset (Biotium) and imaged on a Leica DMi8 microscope (Leica Microsystems). Antibody sources and dilutions are provided in Supplementary Table 2.

Signal intensities in immunofluorescence images were quantified using LasX software (Leica Microsystems) or ImageJ (https://ij.imjoy.io/). For each section, five random areas were chosen, and three non-serial sections per animal were analysed.

#### Analysis of publicly available single cell and bulk RNA-seq data

To assess the clinical relevance of HOXA10 dysregulation, we re-analyzed publicly available bulk and single-cell RNA-seq datasets of human endometrium and decidua (details in Supplementary Table 3). Bulk RNA-seq data from four independent studies included mid-luteal (LH+6 to LH+10) endometrial biopsies from regularly cycling women aged 24–44 years and had not received hormonal treatment for at least two months. Controls had no history of pregnancy loss, RPL cases were defined as women with ≥3 unexplained miscarriages. Raw FASTQ files were subjected to quality assessment using FASTQC, followed by adapter and low-quality read trimming; high-quality reads were aligned to the human genome (GRCh38) using STAR, quantified with GENCODE v45 annotations, and normalized to transcripts per million (TPM) for downstream analyses.

Two single-cell RNA-seq datasets were also examined. The first [7] contained Smart-seq2–derived first-trimester decidual cells. The second dataset [6] included decidua from normal pregnancies (n=5; age 21–35 years; 7–9 weeks gestation) and RPL cases (n=10; at least two prior miscarriages). Both the datasets were processed using Cell Ranger (v9.0.1) and/or Seurat (v5.3.0). From both datasets, *IGFBP6*+ decidual stromal cells were subsetted, followed by clustering, marker identification, and pathway enrichment using WebGestalt. Together, these uniformly processed datasets enabled robust cross-study comparison of HOXA10-associated inflammatory signatures.

#### Statistical analysis

Statistical analyses were performed using GraphPad Prism (version 8). For non-parametric data, comparison between two groups were made using the two tailed Mann-Whitney test. For data where more than two groups were being compared, ANOVA was used followed by Sidak’s post-hoc correction. Correlation analyses were performed using Spearman’s rank correlation method. For all analysis p ≤ 0.05 were accepted as statistically significant.

## Results

### Expression of Hoxa10 is downregulated in the decidua during early pregnancy

Previous studies have shown that HOXA10 expression is downregulated in the decidua of baboons and humans during early pregnancy [26]. Extending these findings in rodents, herein we observed that *Hoxa10* mRNA levels significantly decreased in the mouse endometrium on Days 6 and 7 of pregnancy compared to non-pregnant and Day 5 controls. On Day 8, *Hoxa10* mRNA expression was significantly restored, approaching levels observed on Day 5 (Fig. 1A, n=6 animals/time point).

**Figure 1.**
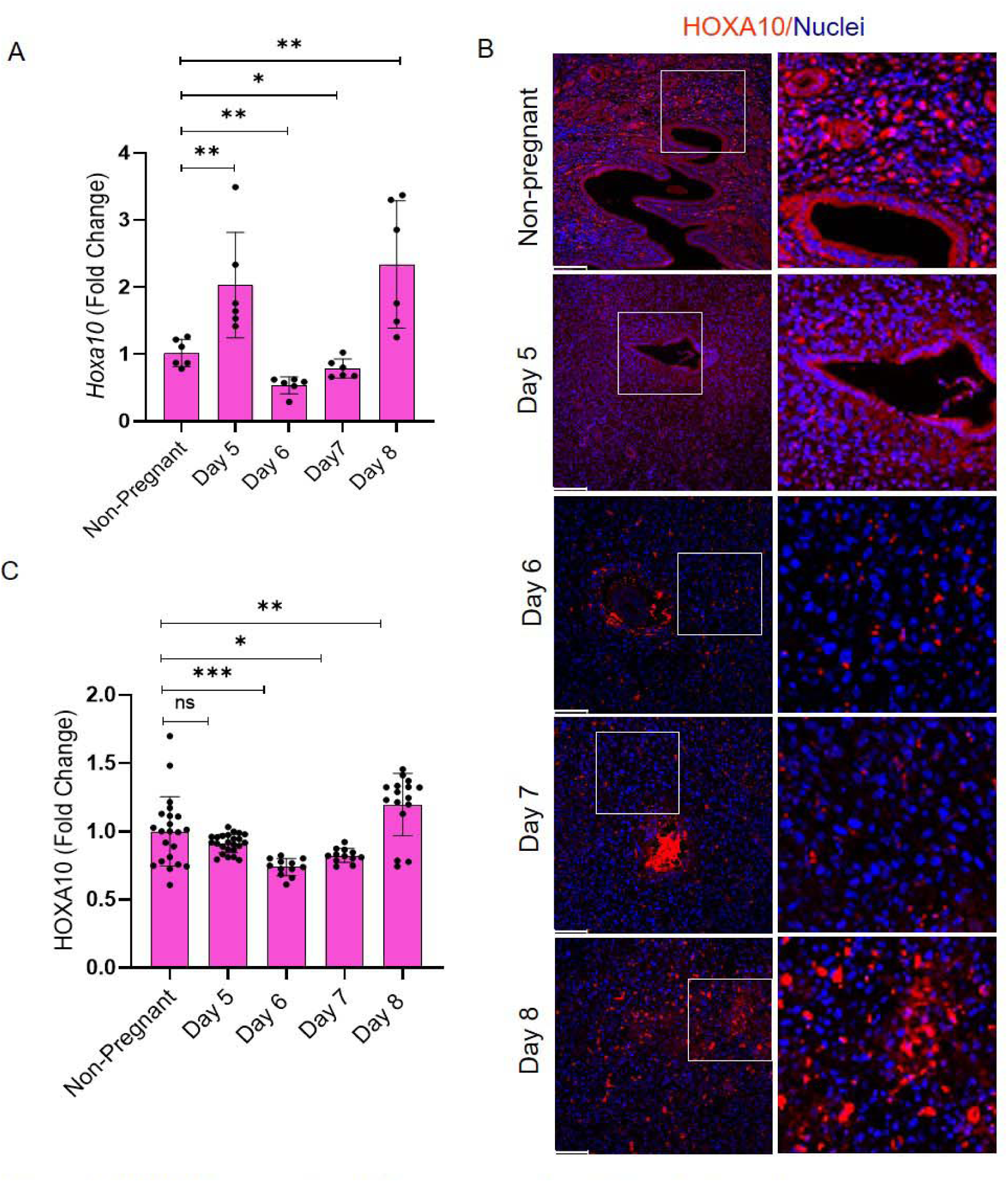
H0XA1O expression in the mouse endometrium during early pregnancy. A) *HoxalO* mRNA levels in the mouse uterus across implantation days (n = 6 animals per time point). Each dot represents one biological replicate, normalized to non-pregnant controls. B) Immunofluorescence for H0XA1O at implantation sites from Day 5-Day 8. Scale bar = 100 µm (zoomed region shown in box). C) Quantification of H0XA1O staining intensity in stromal/decidual cells at the implantation site (n = 3 animals per time point). Data are mean ± SD. ns = not significant, *p < 0.05, **p < 0.01, ***p < 0.001 by two-way ANOVA with Šídäk’s multiple-comparison test.

To assess spatial distribution, we performed immunofluorescence staining for HOXA10. On Day 5 and in non-pregnant controls, intense HOXA10 signals were localized primarily to the stromal compartment. In contrast, on Days 6 and 7, HOXA10 staining was markedly reduced in the decidua. By Day 8, HOXA10 expression was re-established in the decidual zone, including the mesometrial decidua (Fig. 1B, n=3 animals/time point). Quantitative analysis of fluorescence intensity confirmed significant downregulation of HOXA10 on Days 6 and 7, with re-expression on Day 8 (Fig. 1C). No significant changes in HOXA10 expression were observed at inter-implantation sites throughout this period (Supplementary Fig. 1).

These findings indicate that HOXA10 expression is downregulated during the early stages of implantation and restored as pregnancy progresses, suggesting its functional role in the temporal regulation of decidual responses.

### Downregulation of HOXA10 in human decidual stromal cells alters expression of genes in several pathways and increases cell adhesion

Previously we and others have demonstrated that HOXA10 is increased during *in vitro* decidualization of human endometrial stromal cells [27,32]. To explore the functional consequences of HOXA10 downregulation during decidualization, we used siRNA to knock down *HOXA10* in *in vitro* decidualized primary human endometrial stromal cells (hereafter referred to as *HOXA10*KD). Efficient decidualization was confirmed by significant induction of the marker genes *PRL* and *IGFBP1* (Supplementary Fig. 2). Decidualized cells were transfected with *HOXA10* targeting or scrambled control siRNA (scheme described in Fig. 2A). *HOXA10* mRNA levels were reduced by approximately 70% in *HOXA10*KD cells at 24, 48, and 72 hours post-transfection compared to scrambled controls (Fig. 2B, n=5 replicates). To assess the global impact of HOXA10 knockdown, we performed microarrays at all the three time points (Fig. 2A, n=3 replicates per time point). The number of differentially expressed genes (log[FC > 2, p < 0.05) increased over time, with 55 genes altered at 24 h (13 upregulated, 42 downregulated), 604 at 48 h (358 upregulated, 246 downregulated), and 311 at 72 h (192 upregulated, 119 downregulated). Functional enrichment analysis of all the differentially expressed genes revealed significant overrepresentation of biological processes related to cell–substrate adhesion, cytokine signaling, and innate immune responses (Fig. 2C). KEGG pathway analysis showed enrichment of IL17 and TNF signaling pathways, as well as cytokine–cytokine receptor interactions (Fig. 2D).

**Figure 2.**
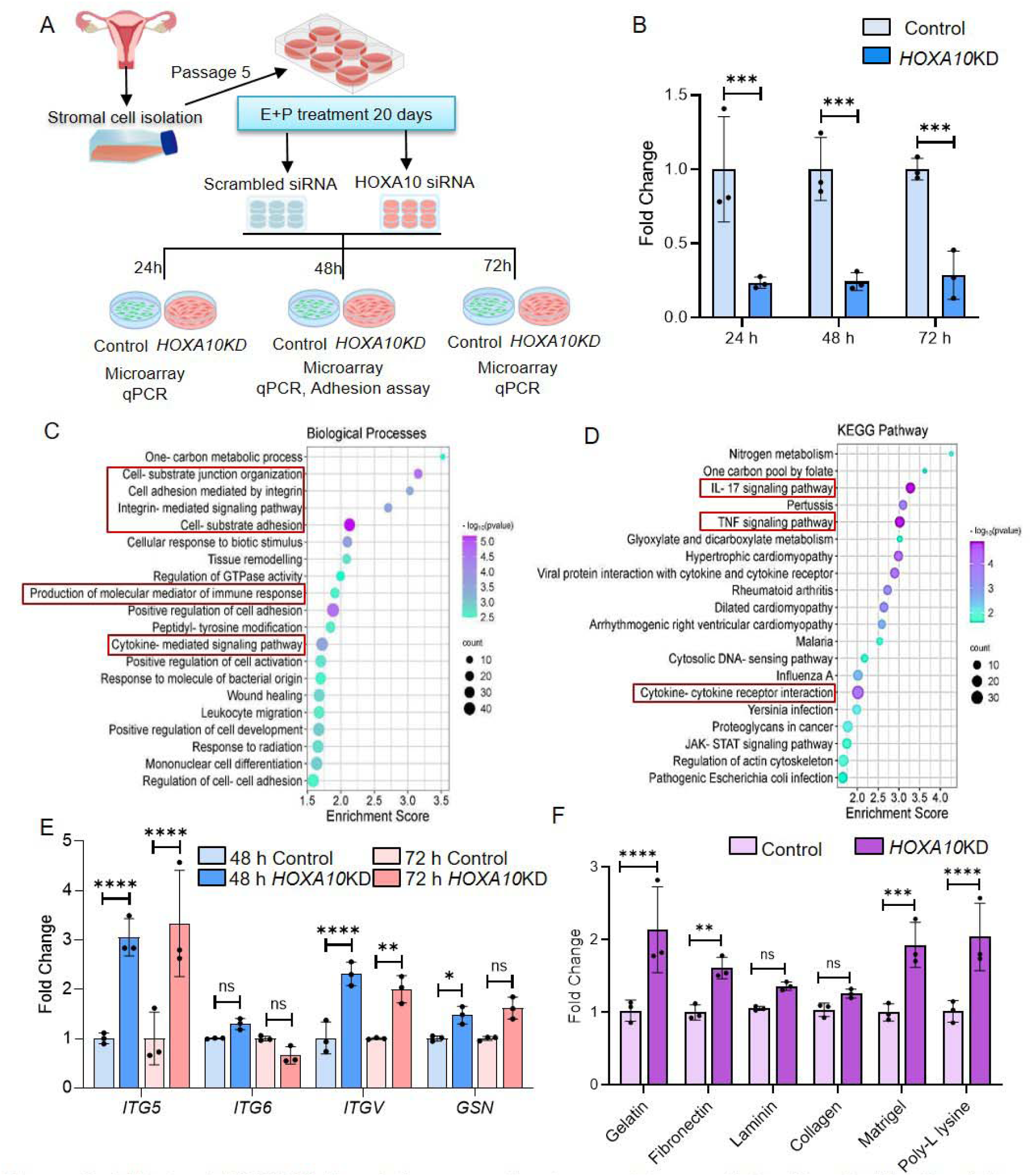
Effect of *HOXA10* knockdown on the transcriptome of *in-vitro* endometrial stromal cells. A) Experimental design. B) *HOXA 10* mRNA levels following siRNA-HOXA10 transfection during *in-vitro* decidualization (n = 3 biological replicates). C-D) Top 20 enriched biological processes (C) and KEGG pathways (D) among differentially expressed genes in HOXA10-knockdown *(HOXA10KD)* cells. **E)** qPCR validation of selected cell-substrate adhesion genes at 48 h and 72 h post-transfection (n = 3). **F)** Adhesion of control and *HOXA* 10KD cells to different substrates (n = 3). Fold change computed relative to mean values of controls (=1). Data are mean± SD. ns = not significant, *p < 0.05, **p < 0.01, ***p < 0.001, ****p < 0.0001 by two-way ANOVA with Sfdak’s correction.

To validate these transcriptomic changes, we measured the expression of key adhesion-related genes (*ITGA5, ITGA6, ITGAV, and GSN*) by qPCR at 48 and 72 h post-transfection. Three of the four genes were significantly upregulated in *HOXA10*KD cells, while the increase in ITGAV did not reach statistical significance (Fig. 2E, n=3 replicates). Functionally, this translated into significantly enhanced adhesion of *HOXA10*KD cells to fibronectin, gelatin, Matrigel, and poly-L-lysine as to compared to scrambled controls (Fig. 2F, n=3 replicates).

These results suggest that HOXA10 regulates the adhesive phenotype of decidual stromal cells, and its downregulation during early decidualization may promote cellular remodeling and matrix interaction, which could be critical for stromal–trophoblast communication at the implantation site.

### Loss of HOXA10 leads to differentiation of inflammatory decidual cells

Along with cell–substrate adhesion, the other striking biological process associated with loss of HOXA10 was cytokine production. To understand this further, we performed STRING–PPI analysis of 56 genes enriched in the biological processes “production of molecular mediators of immune response” and “cytokine-mediated signaling.” Of these, 29 genes formed a single network that contained 10 hub genes (Fig. 3A). Mapping of the 10 hub genes to the top five local network clusters showed that these hub genes were associated mainly with IL1 family signaling and JAK–STAT pathways (Fig. 3B).

**Figure 3.**
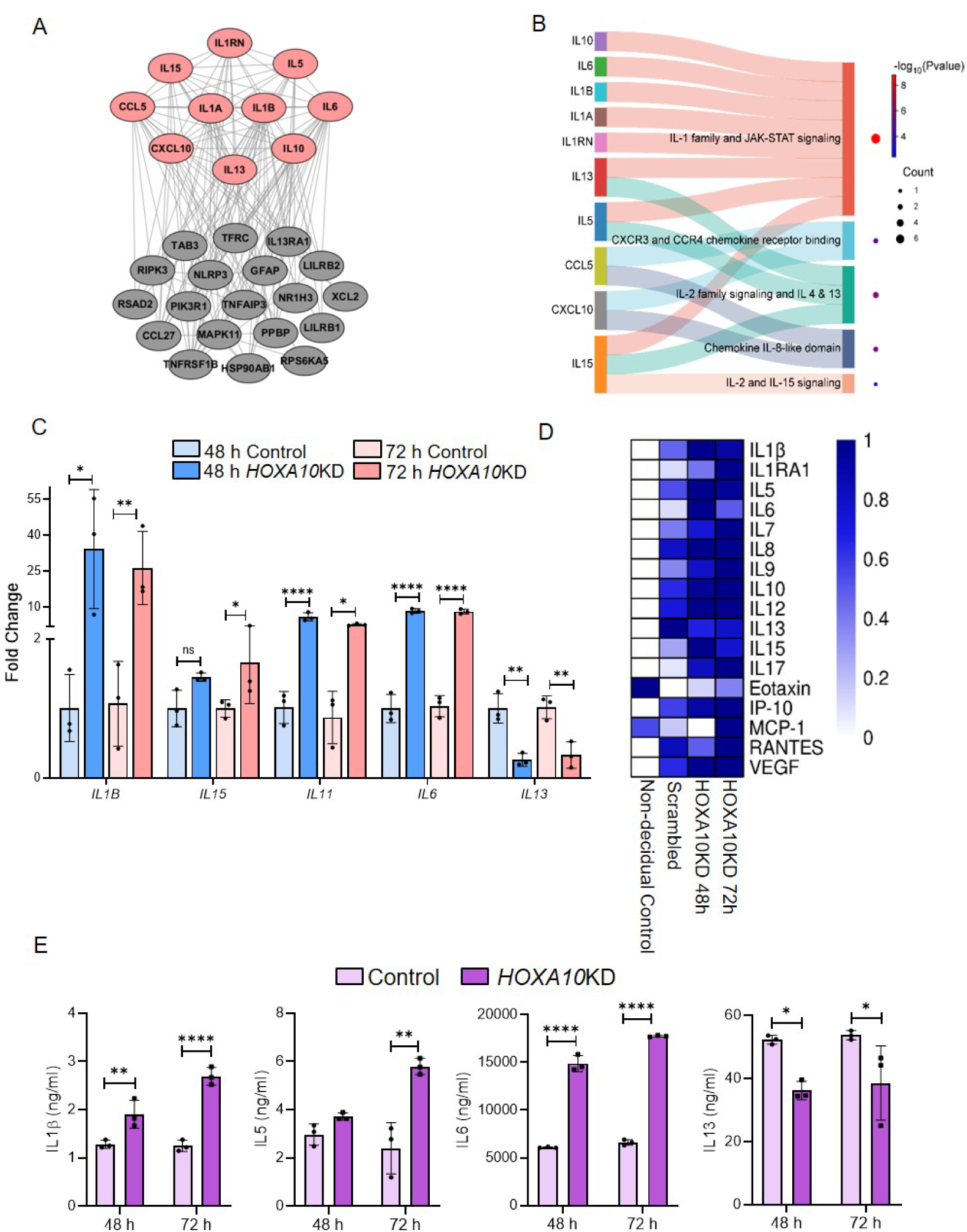
Loss of *HOXA10* promotes the expression of inflammatory markers in decidual cells. A) Protein-protein interaction (PPI) network of differentially expressed genes within the cytokine-cytokine receptor interaction pathway. Hub genes are highlighted in coral. B) Sankey plot showing hub genes and their connections to the top five enriched inflammatory networks. C) qPCR validation of selected hub cytokines in control and *HOXA* 10K□cells at 48 h and 72 h post-transfection (n = 3). D) Heat map of inflammatory cytokines detected in culture supernatants (n = 3 biological replicates) of non-decidualized and decidualized stromal cells transfected with scrambled siRNA (control) or *HOXA10* siRNA *(HOXA* 10KD) at 48 h and 72 h post-transfection. E) Levels of selected hub cytokines in culture supernatants of control and *HOXA* 10KD cells at 48 h and 72 h (n = 3 per time point). Fold change is calculated relative to the mean value of the corresponding control (=1). Data are presented as mean± SD; ns = not significant; *p < 0.05, **p < 0.01, ****p < 0.0001 by two-way ANOVA with Sidak’s multiple-comparison test.

We validated the mRNA levels of five hub genes in *HOXA10*KD cells at 48 and 72 h post-transfection. As compared to control cells, *HOXA10KD* cells had significantly higher expression of *IL1B, IL6, IL11*, and *IL1*5, while *IL13* levels were reduced (Fig. 3C, n=3 replicates). These results prompted us to postulate that loss of HOXA10 may drive an inflammatory phenotype in decidual cells.

To test this, we measured the secretion of 27 inflammation-associated cytokines in the supernatants of scrambled-transfected and *HOXA10*KD decidualized cells, and for comparison, in non-decidualized cells (n=3 replicates per group). Of the 27 cytokines/chemokines/growth factors tested, 17 were above the detection limit of the assay. Compared to non-decidualized controls, several cytokines increased upon decidualization. Downregulation of *HOXA10* further increased the secretion of almost all detectable cytokines, including those corresponding to the hub genes (Fig. 3D and 3E).

At both 48 and 72 h post-transfection, there was a statistically significant increase in secreted levels of the hub cytokines IL1β, IL5, and IL6, while IL13 levels were significantly lower in *HOXA10*KD cells compared to controls (Fig. 3E, n=3 replicates). Together, these results suggest that loss of HOXA10 in decidualized human endometrial stromal cells increases the expression and secretion of inflammatory cytokines, and therefore we termed these as inflammatory decidual stromal cells (iDSCs).

### Emergence of inflammatory decidual stromal cells marked by IL1**β** during implantation in the mouse uterus

Having established that HOXA10 downregulation *in vitro* induces inflammatory cytokines and the differentiation of iDSCs, we next examined whether iDSCs emerge during the peri-implantation window in a manner consistent with HOXA10 regulation. To address this, we assessed the expression of IL1β, the hub inflammatory cytokine induced upon HOXA10 loss, in mouse uteri from Days 5 to 8 of pregnancy (n=3 animals per time point).

Immunofluorescence revealed low levels of IL1β on Day 5, prior to full decidualization, its expression increased substantially on Days 6 and 7, coinciding with the active implantation phase (Fig. 4A). By Day 8, IL1β expression had significantly declined. Quantitative analysis confirmed this dynamic pattern (Fig. 4B). During this same period, HOXA10 levels decreased on Days 6 and 7 and increased again on Day 8 (Fig. 4C).

**Figure 4.**
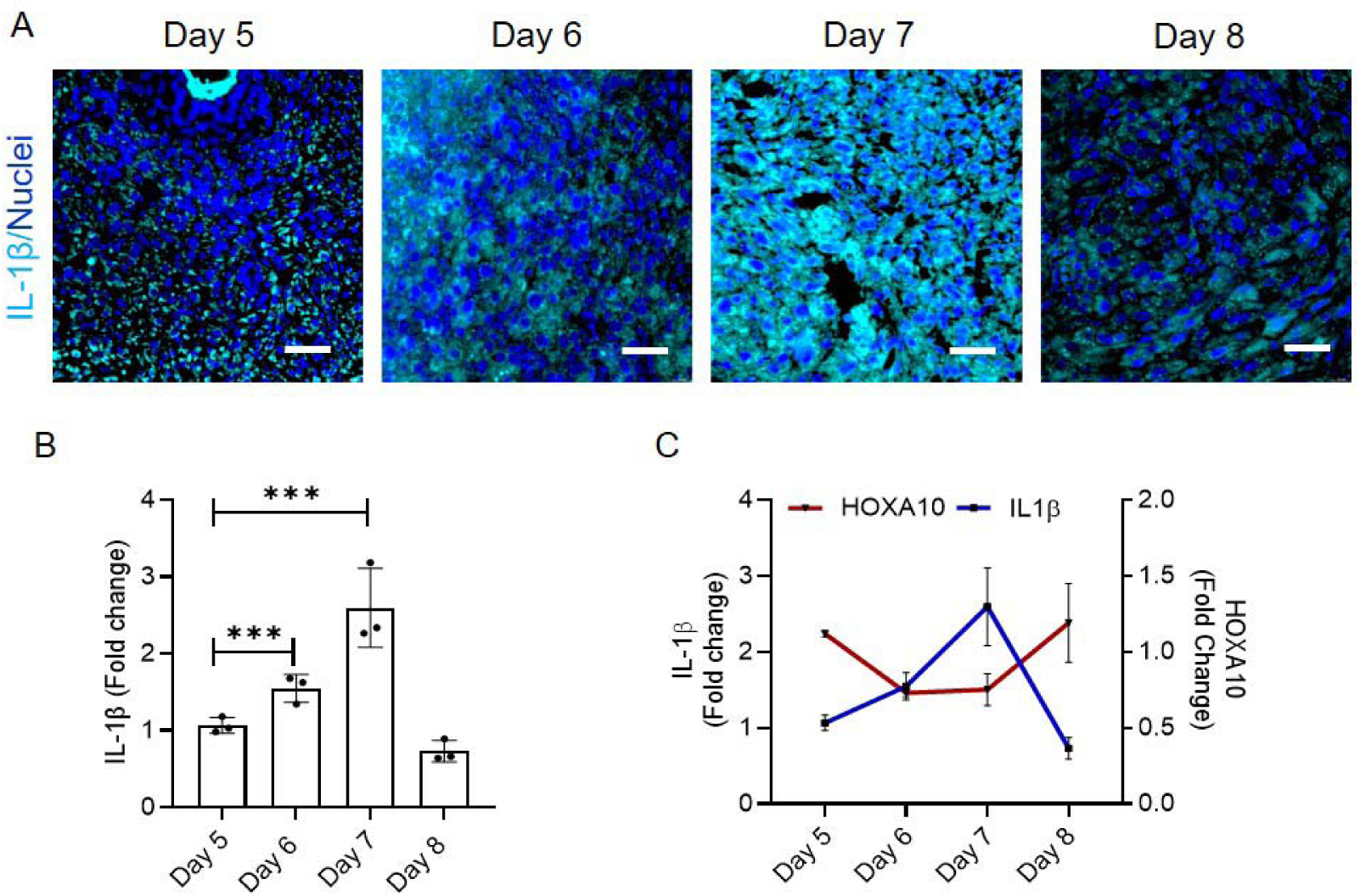
Reciprocal expression of IL1 β and H0XA1O in the mouse endometrium during implantation. A) Immunofluorescence staining for IL1 β at implantation sites from Day 5 to Day 8 (n = 3 animals per time point). Scale bar = 50 µm. B) Quantification of IL1 β staining intensity across implantation days. C) Quantification of stromal/decidual H0XA1O and IL1 β levels across implantation days, fold change relative to Day 5 (=1). Data are mean *SĐ. ***p <0.001 by two-way ANOVA with Šídák’s test.

These findings support the model that the appearance of iDSCs during implantation is coordinated with the transient downregulation of HOXA10 (Fig. 1), and that re-expression of HOXA10 is likely required for the resolution of this inflammatory phenotype.

### HOXA10 is low in the iDSCs of the first trimester human decidua

To determine whether HOXA10 is associated with iDSCs in human decidua, we analyzed scRNA-seq data from first-trimester human decidua [7]. *IGFBP6*+ cells were annotated as decidual stromal cells [6,7], and high-quality data from 375 cells were projected onto a UMAP for dimensional reduction. As reported previously, decidual stromal cells segregated into three clusters (ds1, ds2, and ds3). *HOXA10* expression was lowest in the ds2 cluster relative to ds1 and ds3 (Fig. 5A). Differential gene expression followed by GO enrichment analysis confirmed the identity of these clusters. The ds1 cluster was enriched for biological processes related to muscle development, consistent with myofibroblastic decidual stromal cells. The ds2 cluster showed enrichment for inflammatory pathways, matching the previously described iDSC phenotype [6,7]. The ds3 cluster was enriched for stress-response pathways (Fig. 5B). Notably, the ds2 cluster, which showed the lowest HOXA10 expression, exhibited higher expression of IL1A, IL1B, IL1R2, and other inflammatory mediators (Fig. 5C–E). These findings indicate that, similar to the mouse, low HOXA10 expression in human decidua is associated with the inflammatory decidual stromal cell (iDSC) state.

**Figure 5.**
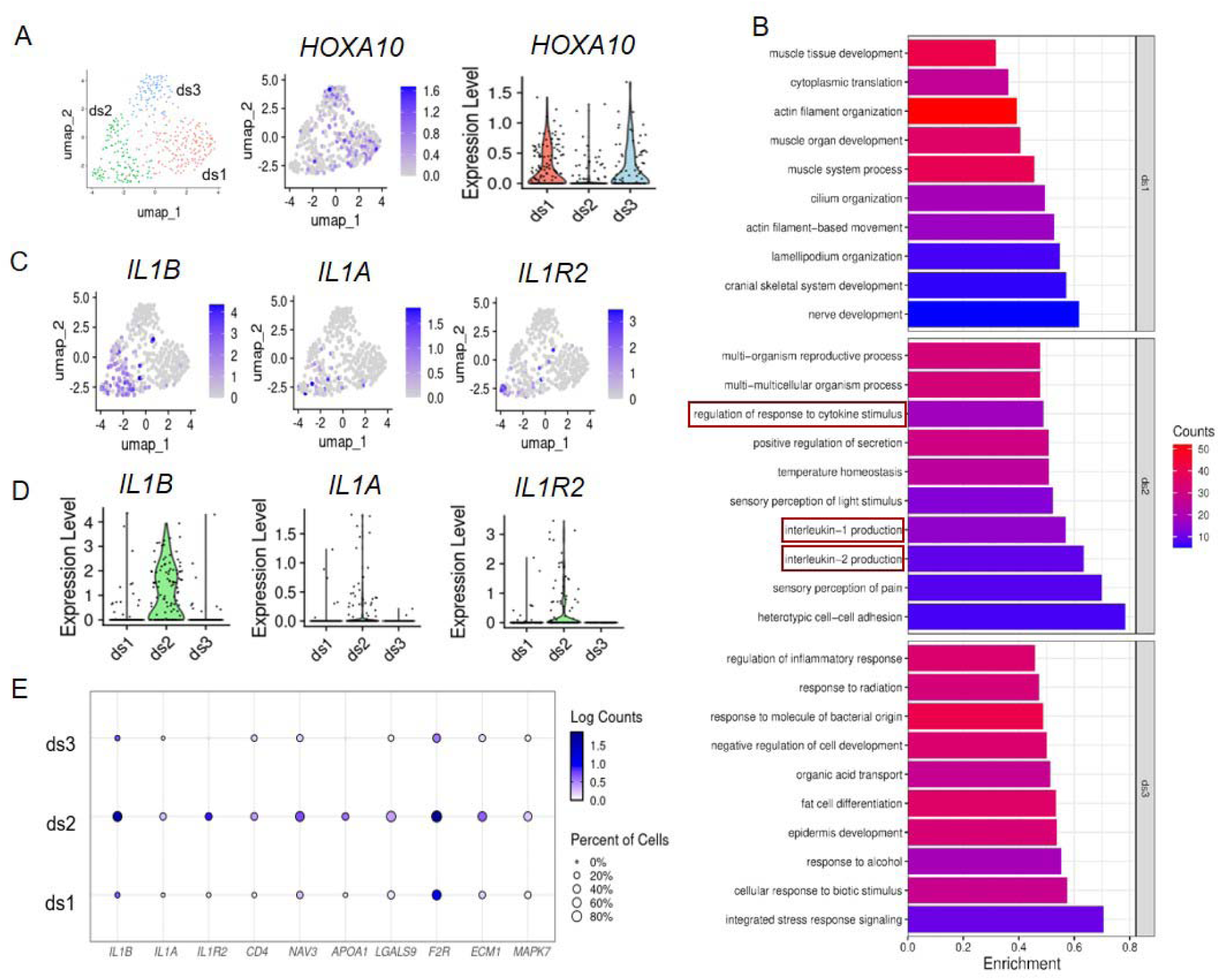
*H0XA1O* is fow in inflammatory decidual cell clusters in first-trimester human decidua. A) UMAP of 375 decidual stromal cells (ENA: E-MTAB-6678), showing *H0XA1O* expression across clusters dsl, ds2, and ds3. B) Enriched biological processes in differentially expressed genes for each cluster. C) Feature plots for selected genes. D) Violin plots showing expression of *IL1B, IL1A,* and *IL1R2* across clusters. E) Dot plot showing expression of genes related to enriched biological processes associated with cytokine production.

### Loss of HOXA10 induces inflammation and inflammasome activation in the non-pregnant endometrial stroma

To determine whether loss of HOXA10 activates inflammation in the endometrial stroma *in vivo*, we analyzed uteri from non-pregnant *Hoxa10* hypomorphic mice that has reduced HOXA10 levels (Fig. 6A, Supplementary Fig. 3A). Quantitative RT-PCR of endometrial tissue in the diestrus phase revealed significantly elevated expression of the inflammatory cytokines *Il1b, Ifng*, and *Il*6 in hypomorphs (n=3) compared to wild-type controls (Supplementary Fig. 3B). These findings suggest that HOXA10 deficiency is sufficient to induce a basal pro-inflammatory state in the endometrium even in the absence of decidualization.

**Figure 6.**
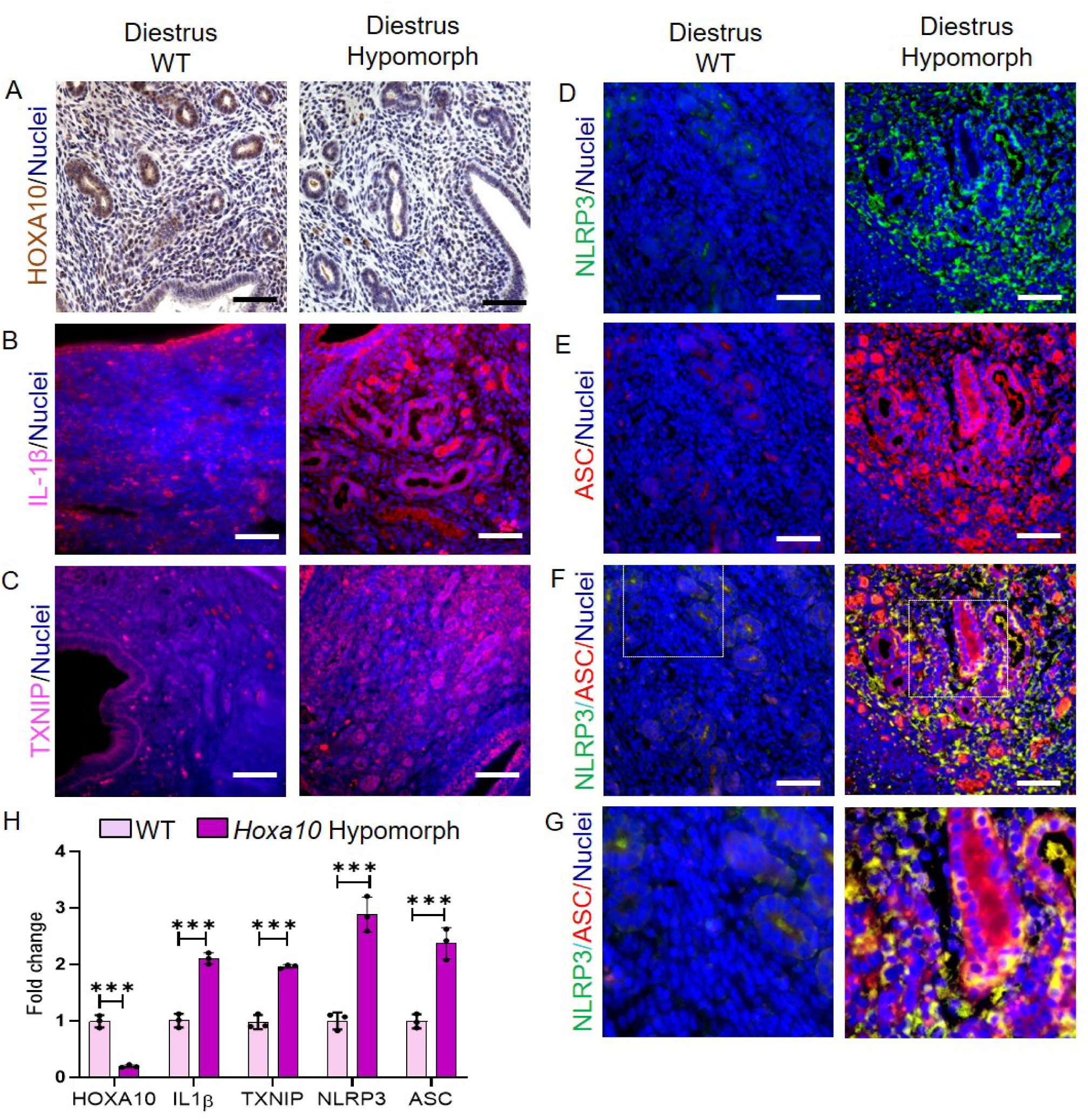
Loss of HOXA10 drives an inflammatory phenotype in endometrial stromal cells. A-E) Immunolocalization of HOXA10 (A), IL1β (B), TXNIP (C), NLRP3 (D), and ASC (E) in wild-type (WT) and *Ho×alO* hypomorph mice at the diestrus-stage (n = 3 animals per group). F) Co-localization of NLRP3 and ASC in the endometrium. G) Zoomed view of the boxed region in panel F. Scale bar = 50 µm. H) Quantification of immunostaining intensity, where each dot represents the mean value from one animal (n = 3 biological replicates). Y-axis values represent fold change, calculated by taking the mean intensity of WT controls as 1. Data are presented as mean ± SD; ***p < 0.001, determined by two-way ANOVA with Šídák’s multiple-comparison test.

To localize this inflammatory response, we performed immunofluorescence staining for TXNIP and IL1β (the markers of iDSCs). In wild-type animals (n=3), both proteins were minimally detected in stromal cells. In contrast, stromal cells of hypomorphs showed robust expression of both IL1β and TXNIP (n=3; Fig. 6B and C). Because IL1β secretion requires inflammasome activation, we next examined the expression and localization of the core inflammasome components NLRP3 and ASC. Both proteins were barely detectable in the stromal cells of wild-type mice (n=3). In hypomorphs (n=3), however, NLRP3 and ASC were abundantly expressed and formed prominent punctate foci (Fig. 6D and E). Dual immunofluorescence (n=3 for both controls and hypomorphs) revealed strong colocalization of NLRP3 and ASC in hypomorph stromal cells, indicating inflammasome complex formation (Fig. 6F and G). Quantitative analysis confirmed a significant decrease in HOXA10 and an increase in the abundance of all inflammasome-associated proteins in HOXA10-deficient stroma (Fig. 6H).

Together, these findings demonstrate that loss of HOXA10 in the non-pregnant uterus is sufficient to trigger a pro-inflammatory response and promote inflammasome assembly in endometrial stromal cells, independent of decidualization.

### Persistent inflammatory decidual stromal cells in pregnant HOXA10 hypomorphs

We next asked whether persistent HOXA10 deficiency post-implantation affects the resolution of decidual inflammation. We examined uteri from pregnant *Hoxa10* hypomorphs (n=3 animals, 3 implantation sites each) and wild-type controls (n=3 animals, 3 implantation sites each) on Day 9 of pregnancy.

In wild-type mice, HOXA10 expression was robustly restored in the decidua by Day 9, and both IL1β and TXNIP markers of iDSCs were minimally detectable (Fig. 7A–C). In contrast, Hoxa10 hypomorphs exhibited low HOXA10 levels and had strong expression of both IL1β and TXNIP in the decidua (Fig. 7A–C). To determine whether inflammasome activity also persisted, we assessed the expression and colocalization of NLRP3 and ASC. In wild-type decidua, these proteins were barely detectable (Fig. 7D and E), and only a few cells showed colocalization (Fig. 7F and G). In contrast, the decidua of HOXA10 hypomorphs contained numerous cells with strong NLRP3 and ASC expression, along with frequent colocalized puncta indicating continued inflammasome assembly (Fig. 7D–G). Quantitative analysis revealed a significant reduction in HOXA10 and a corresponding increase in IL1β, TXNIP, NLRP3, and ASC in hypomorphs compared to controls (Fig. 7H).

**Figure 7.**
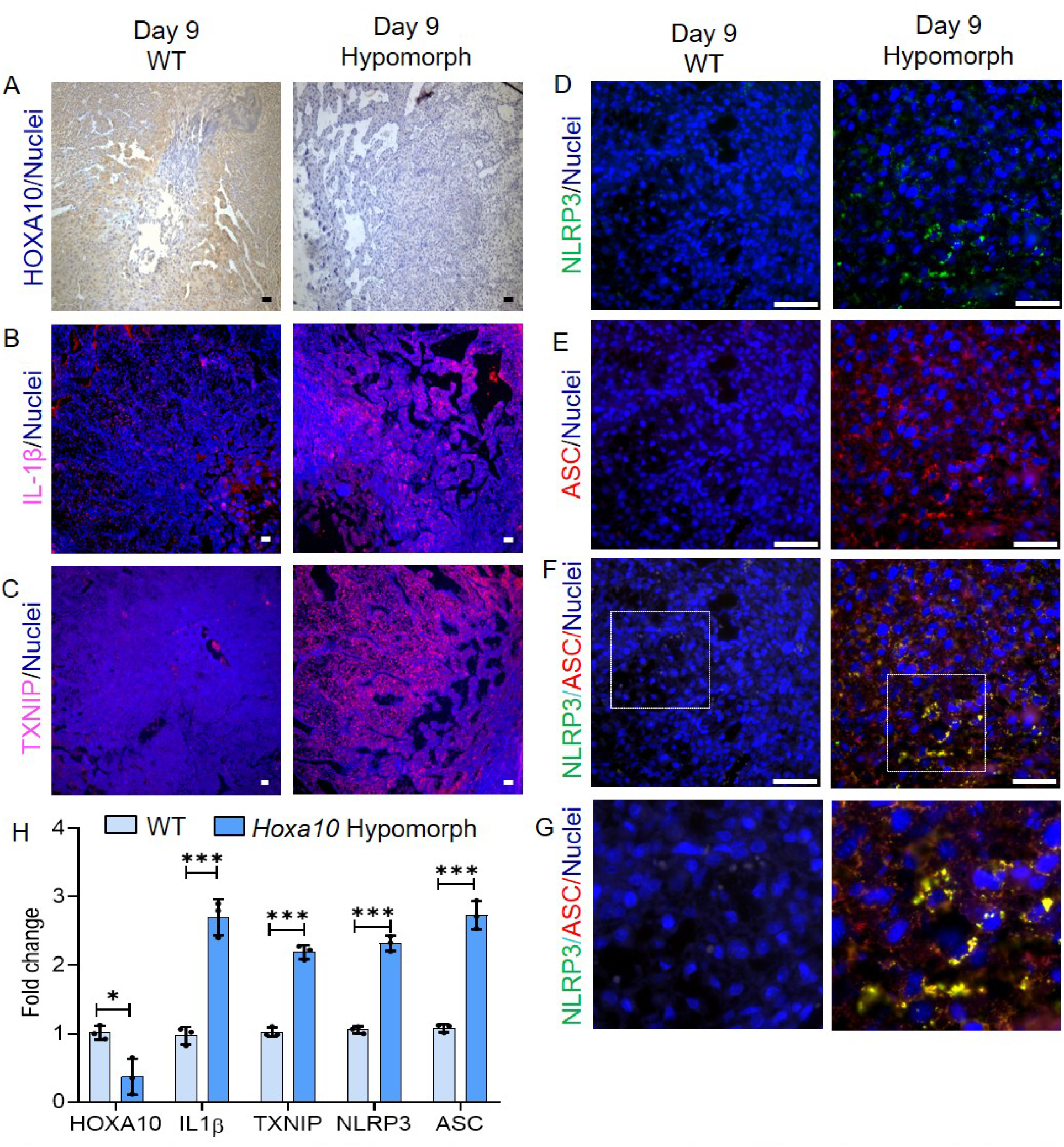
Inflammatory decidual cells persist in the endometrium of pregnant *Hoxa10* hypomorphs. A-E) lmmunolocalization of HOXA10 (A), IL113(B), TXNIP (C), NLRP3 (D), and ASC (E) in wild-type (WT) and *Hoxa10* hypomorph mice on day 9 of pregnancy (n = 3 animals per group). F) Co-localization of NLRP3 and ASC in the endometrium. G) Zoomed view of the boxed region in panel F. Scale bar = 50 µm. H) Quantification of immunostaining intensity, where each dot represents the mean value from one animal (n = 3 biological replicates). Y-axis values represent fold change, calculated by taking the mean intensity of WT controls as 1. Data are presented as mean ± SD; *p < 0.05, ***p < 0.001, determined by two-way ANOVA with Sidak’s multiple-comparison test.

These findings suggest that restoration of HOXA10 after implantation is necessary to suppress iDSC activity. Its absence leads to the persistence of an inflammatory decidual phenotype marked by elevated IL1β, TXNIP, and active inflammasomes an environment likely detrimental to pregnancy maintenance.

### HOXA10 deficiency leads to subfertility, implantation defects, and abnormal placental development

To determine the physiological consequences of persistent decidual inflammation in *Hoxa10* hypomorphs, we evaluated fertility, implantation, and placental architecture in comparison to wild-type controls.

Assessment of reproductive performance revealed that hypomorphs had a significantly reduced mean litter size compared to wild-type animals (n=10/group; Fig. 8A). To assess reproductive capacity over time, serial mating experiments were performed (scheme explained in Fig. 8B; n=20 hypomorphs). While 15 of 20 hypomorphs conceived during the first cycle, only 7 conceived in the second, and 2 in the third. None conceived by the fourth cycle, indicating progressive loss of fertility (Fig. 8C). Moreover, even among animals that initially conceived, litter size decreased with each subsequent pregnancy.

**Figure 8.**
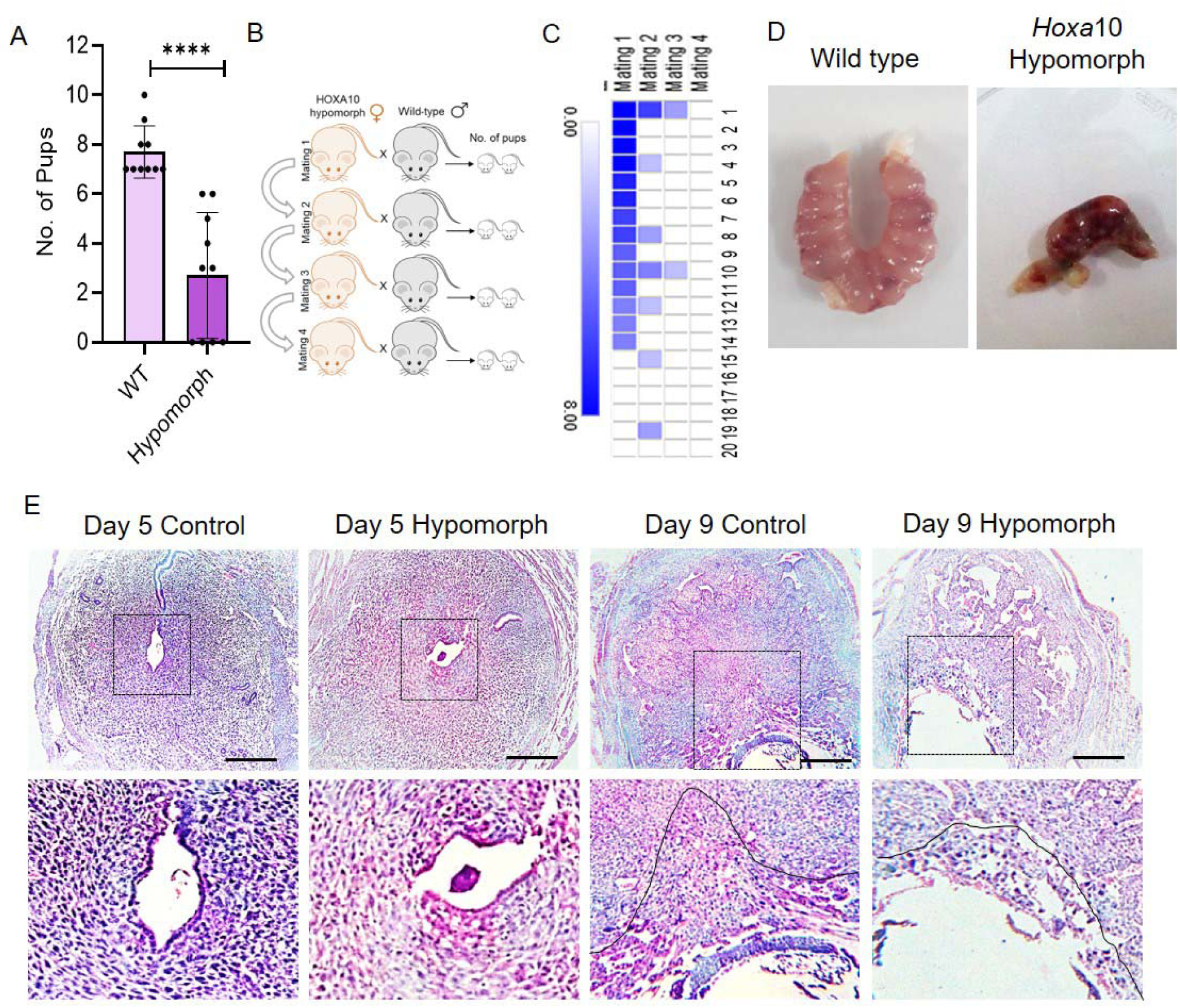
Subfertility and defective placentation tn *Hoxa10* hypomorphs. A) Litter size in *Hoxa10* hypomorphs compared with wild-type controls (n = 10 animals per group). B) Experimental strategy for the serial mating paradigm. C) Heat map showing the number of pups born per mating in *Hoxa10* hypomorphs (π = 20 animals). D) Representative images of uteri from wild-type and *Hoxa10* hypomorph mice on Day 9 of pregnancy. E) Hematoxylin- and eosin-stained endometrium from wild-type and *Hoxa10* hypomorphs on Day 5 and Day 9 of pregnancy. Scale bar = 50 µm (boxed region shows a higher-magnification view). The placentation area is indicated. Data are presented as mean ± SD. “**p < 0.0001 by two-tailed Mann-Whitney test.

Gross examination of Day 9 pregnant uteri (n=3 animals) revealed fewer implantations in *Hoxa10* hypomorphs (Fig. 8D). Histological analysis further confirmed that implantation sites in hypomorphs exhibited disorganized architecture. On Day 5 (n=3 animals/group), many embryos in hypomorphs failed to properly attach to the endometrium (Fig. 8E). On Day 9 (n=3 animals, 4 implantation sites each/group), the wild-type implantation sites displayed well-formed ectoplacental cones and organized labyrinthine trophoblasts, hypomorphs showed flattened, disorganized placental regions lacking a clear ectoplacental structure (Fig. 8E).

Together, these findings indicate that persistent decidual inflammation driven by HOXA10 deficiency leads to compromised implantation, impaired placental morphogenesis, and progressive subfertility.

### Women with recurrent pregnancy loss have reduced *HOXA10* and elevated inflammatory gene expression in the endometrium and decidua

To assess the clinical relevance of our findings, we analyzed bulk RNA-seq data from mid-luteal endometrial biopsies of women and those with recurrent pregnancy loss (RPL). High-quality data of 30 fertile controls and 17 RPL patients were analyzed. *HOXA10* mRNA levels were significantly lower in RPL samples, whereas *IL1B* and *PYCARD* were significantly elevated (Fig. 9A), indicating a heightened inflammatory state.

**Figure 9.**
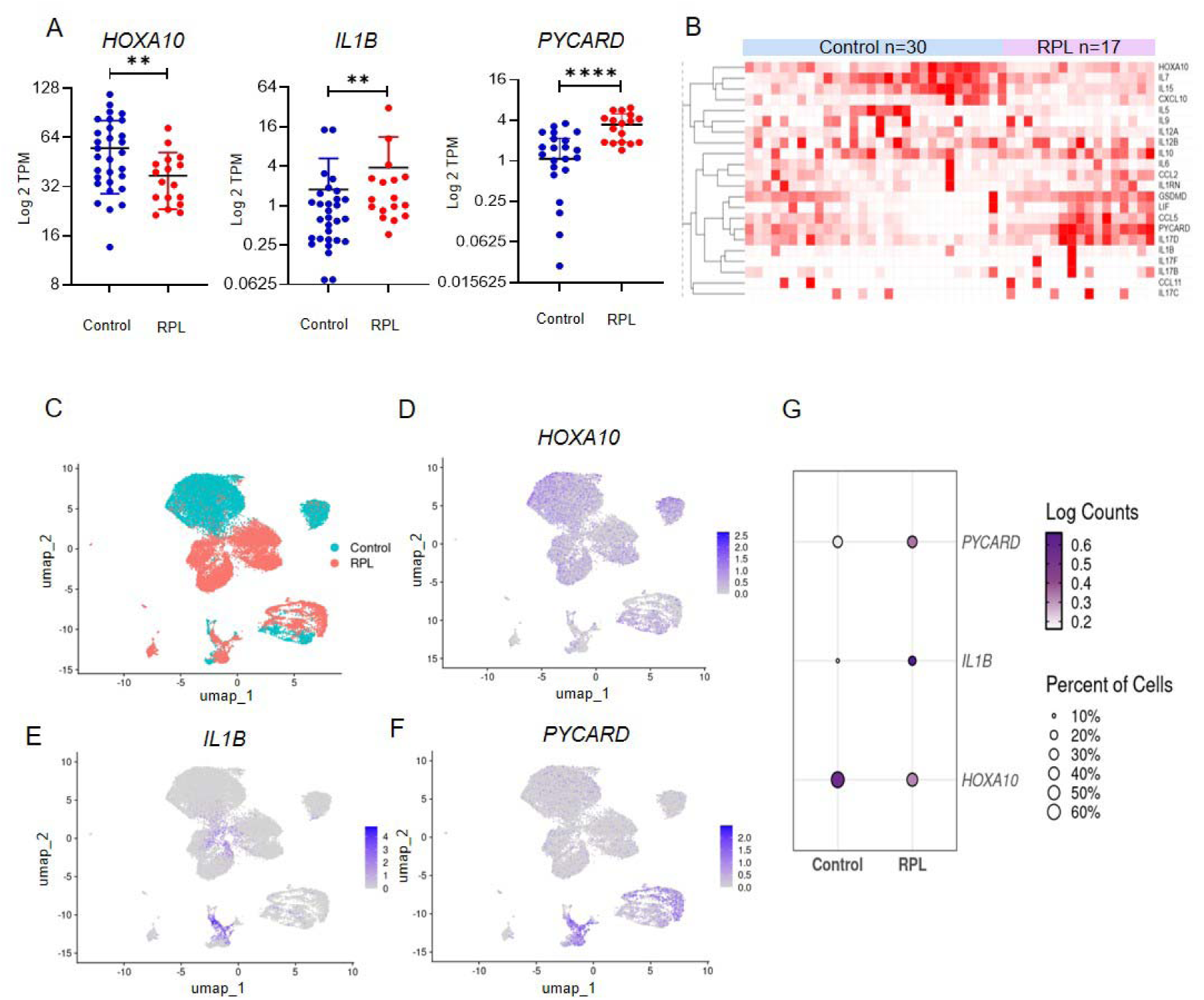
Recurrent Pregnancy Loss is associated with low *HOXA10* and high is inflammatory gene expression in the human endometrium and first-trimester decidua. A) TPM levels of *HOXA10, 1L1B,* and *PYCARD* in mid-luteal endometrial biopsies from controls dow (π = 30) and RPL patients (n = 17), aggregated from four independent publicly available datasets. Each dot represents one patient. Data are mean ± SD. p-values from two-tailed Mann-Whitney test (“p < 0.01; “**p < 0.0001). B) Heat map of selected *HOXA1* O-regulated inflammatory genes in control vs RPL endometrium. C) UMAP of *IGFBP6+* decidual cells from 22 first trimester decidua (OEP0Ũ29Ũ1) in control vs RPL cases. Feature plots showing expression of D) *HOXA10,* E) *1L1B,* and F) *PYCARD* in decidual cells. G) Dot plot summarizing percentage of expressing cells and expression intensity of the three genes.

We next examined 20 inflammatory genes that were upregulated in *HOXA10*KD decidual stromal cells. Nearly half of the RPL samples showed coordinated upregulation of multiple inflammatory mediators compared to controls (Fig. 9B).

To determine whether these inflammatory changes occur specifically in decidual stromal cells, we analyzed *IGFBP6+* stromal cells from a first-trimester scRNA-seq dataset (NODE: OEP002901). High-quality profiles were obtained from 12,821 cells from controls and 13,126 cells from RPL cases. *HOXA10* expression was lower in decidual stromal cells from RPL samples, while *IL1B* and *PYCARD* were markedly higher (Fig. 9C–D). Across both bulk and single-cell datasets, *HOXA10* levels showed a modest but statistically significant negative correlation with *IL1B* and *PYCARD* expression (Supplementary Fig. 5).

Together, these findings indicate that reduced HOXA10 expression and aberrant activation of inflammatory pathways in decidual stromal cells are associated with RPL, linking our *in vitro* and mouse data to human pregnancy loss.

## Discussion

The results of this stud y dem onst rate that HO XA 10 is dow nregulated in the decidua around the time of implantation, which leads to the differentiation of decidual cells with an inflammatory phenotype. An increase in HOXA10 expression post-implantation is required to resolve this inflammation. If HOXA10 remains low beyond implantation, it results in persistent inflammation that is associated with defective placentation and subfertility. In the human endometrium, low HOXA10 and elevated inflammatory markers are associated with RPL.

Pregnancy in mammals involves the immunological acceptance of a semi-allogeneic fetus and is typically associated with an anti-inflammatory maternal environment. Paradoxically, implantation initiates an inflammatory cascade in the maternal endometrium, that must be suppressed soon after for pregnancy to progress [8,9,12,14,15,26,33]. However, the molecular regulators that orchestrate this temporal switch in inflammation are poorly understood. Our data identify endometrial HOXA10 as one such regulator. Contrary to the long-held belief that HOXA10 expression is uniformly required during pregnancy, our earlier work showed that HOXA10 is downregulated in the early decidua of baboons and humans [26]. Extending those findings, we now show that HOXA10 is similarly reduced in the decidualizing stroma of mice during implantation, confirming this as a conserved phenomenon.

To understand the function of HOXA10 loss in decidual stromal cells, we silenced its expression in primary human stromal cells undergoing in vitro decidualization and observed the enrichment of pathways involved in actin cytoskeleton organization and adhesion, accompanied by changes in integrin expression and enhanced adhesion to extracellular matrices. Previous studies that performed global expression profiling of endometrial stromal cells following HOXA10 knockdown also showed several genes associated with actin remodeling and integrins [32,34]. HOXA10 is known to bind the promoter of β3-integrin, and regulates its expression in endometrial cells [35]. These results suggest that HOXA10 would modulate the adhesive and structural phenotype of decidual cells during implantation. Integrin switching is a well-reported feature of decidual cells during implantation and is considered important for trophoblast invasion in multiple species [36–42]. Taken together, these studies imply that HOXA10 in decidualizing stromal cells controls the expression of genes involved in cell adhesion, which may be required for physiological functions associated with implantation.

Along with changes in adhesion, another striking consequence of HOXA10 downregulation was the induction of inflammation-related genes. The differentially expressed genes were enriched in IL1 family and JAK–STAT signaling pathways. HOXA10-silenced cells secreted increased levels of IL1β, IL6, IL11, IL15, and several chemokines. Protein–protein interaction analysis identified a cluster of 10 hub cytokines associated with inflammation, most of which were significantly upregulated at both transcript and protein levels. Recent scRNA-seq studies of first-trimester human decidua identified three major decidual stromal subtypes: an inflammatory subtype termed inflammatory decidual stromal cells (iDSCs), a glycolytic subtype, and a myofibroblastic subtype [6,7]. Additionally, scRNA-seq of human stromal cells undergoing *in vitro* decidualization has shown upregulation of ER stress and inflammatory markers, including TXNIP and IL1B, in a subset of cells [43,44]. Corroborating with these studies we did identify the inflammatory decidual stromal cells (iDSCs) and the myofibroblastic subtypes in the first trimester decidua and and notably, *HOXA10* expression was lowest in the iDSCs, suggesting that reduced HOXA10 is associated with iDSC differentiation. Since HOXA10 is reduced during the early decidual response, and its downregulation in vitro promotes inflammatory cytokine production, it is plausible that HOXA10 loss may act as a trigger for iDSC emergence.

During implantation, several studies have reported increased NLRP3 expression in the endometrium, suggesting activation of inflammatory pathways during implantation [43,45–48]. To test whether HOXA10 controls inflammation in vivo, we examined the endometrium of non-pregnant HOXA10 hypomorphic mice. Even in the absence of decidualization, these mice showed elevated expression of IL1β and TXNIP in stromal cells, along with localized puncta of NLRP3 and ASC, indicating inflammasome activation. These observations suggest that HOXA10 suppresses basal inflammatory signaling in the endometrium and that its loss leads to inflammasome priming. This is particularly relevant for endometrial disorders such as endometriosis, where HOXA10 is known to be reduced [19,22,24,30,49–51] and inflammation-related factors, including IL1β and inflammasome components, are elevated [46,52–54]. In the context of infertility, a subset of hypomorphs were infertile, and most animals showed disrupted expression of receptivity markers. Reduced HOXA10 is also reported in the endometrium of women with unexplained infertility and recurrent implantation failure [19,51]. Thus, our data suggest that a transient reduction of HOXA10 contributes to the inflammatory burst required at implantation, whereas premature or persistent reduction is associated with pathological inflammation associated with infertility.

We observed that post-implantation, HOXA10 levels increase, coinciding with a reduction in inflammation in vivo. We next asked whether this rise in HOXA10 is essential to extinguish post-implantation inflammation. In pregnant hypomorphs, Day 9 decidua retained high levels of IL1β and TXNIP, and active NLRP3–ASC inflammasomes were abundant. These findings indicate that HOXA10 re-expression is necessary to suppress inflammation after implantation. In its absence, the decidua remains inflamed. Interestingly, in the hypomorphs that did become pregnant, implantation sites showed disorganized ectoplacental structures, suggesting that resolution of inflammation is essential for proper placentation. This is consistent with previous reports demonstrating that inflammatory cytokines, including many of the hub cytokines identified here, regulate trophoblast invasion [11,12,26,55–57]. Furthermore, in humans, excessive decidual inflammation is associated with shallow trophoblast invasion and preeclampsia [11,45,58,59] These findings support the conclusion that HOXA10 is the switch required for the timely resolution of inflammation post-implantation an essential step for proper placentation.

RPL is defined as the loss of two or more pregnancies before 24 weeks of gestation, and in nearly 50% of cases the cause is unknown. Multiple risk factors have been associated with RPL, including genetic abnormalities, endocrine dysfunction, paternal factors, the uterine microbiome, and abnormal decidual cellular function [3,5,6,60,61]. A growing body of evidence suggests that many of these factors converge on endometrial dysfunction and immune dysregulation including inflammasome activation [6,18,62–65]. To assess the relevance of our findings in this context, we analyzed endometrial bulk and decidual scRNA-seq data from women with RPL. *HOXA10* expression was significantly lower in RPL samples, while *IL1B* and *PYCARD* were upregulated. In a subset of RPL cases with low *HOXA10*, several inflammatory genes identified in *HOXA10*KD cells were also elevated. Our reanalysis of scRNA-seq data confirmed that *HOXA10* is reduced in decidual stromal cells from RPL cases and that this reduction is accompanied by increased inflammasome marker expression. Together, these findings support a model in which loss of HOXA10 promotes decidual inflammation, and failure to resolve this response is associated with miscarriage. In light of these findings, it will be important to determine how RPL risk factors disrupt the HOXA10–inflammasome axis and influence iDSC differentiation.

Finally, how HOXA10 regulates these inflammatory programs remains an open question. As a transcription factor, HOXA10 may cooperate with other regulators such as STAT family proteins or CTCF. Recent cistromic analyses show that HOXA10 binding sites overlap with CTCF and STAT motifs [25]. It is likely that similar interactions occur in stromal cells to regulate long-range chromatin loops controlling inflammatory gene expression. Mapping HOXA10 occupancy in decidual stromal cells will help uncover this mechanism.

In summary, we propose that loss of HOXA10 initiates a transient inflammatory phenotype in decidual stromal cells that facilitates implantation, and HOXA10 re-expression is then required to suppress this inflammation and support placentation. Disruption of this temporal regulation is associated with reproductive failure in mice. In humans, reduced HOXA10 correlates with increased inflammatory gene expression in the decidua of women with RPL, supporting its role in pregnancy maintenance.

## Data availability statement

The microarray data is available at the GEO datasets with the accession ID GSE35801. The following datasets PRJNA273001, PRJNA314429, PRJNA342633, PRJNA384963, PRJEB25794, OEP002901 were reanalyzed in the present study.

## Acknowledgements

The study bears the NIRRCH ID: RA/1914/07-2025. The support of ICMR intramural grants is acknowledged. RS is a recipient of the DST Inspire fellowship. SS is a recipient of the DBT Biocare Women Scientist. RP is the recipient of CSIR fellowship and AcSIR PhD student. We declare that ChatGPT Go -5.1, under the supervision and input of the authors was used for language editing this manuscript.

## Authors Role

RS and DM conceptualized the idea, RS, BN, AM, and GG performed the experiments. RS, BN, SS, SP, GG, AM, and PR did data analysis. DM, BN, and RS drafted the paper. All the authors critically revised the paper and accepted the final version of the manuscript. All authors agree to be accountable for all aspects of the work.

