## Supplementary figures and images for "Temporal Control of Decidual Inflammation by HOXA10 is Essential for Implantation and its Dysregulation is Associated with Early Pregnancy Loss"

### Supplymental Fig 1

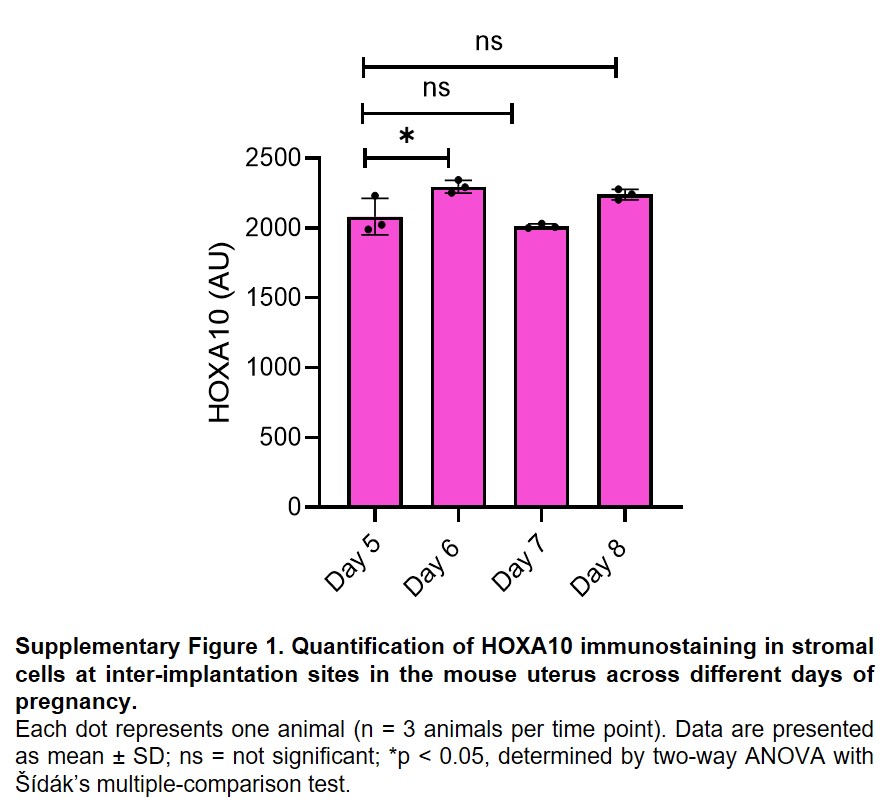

### Supplymental Fig 2

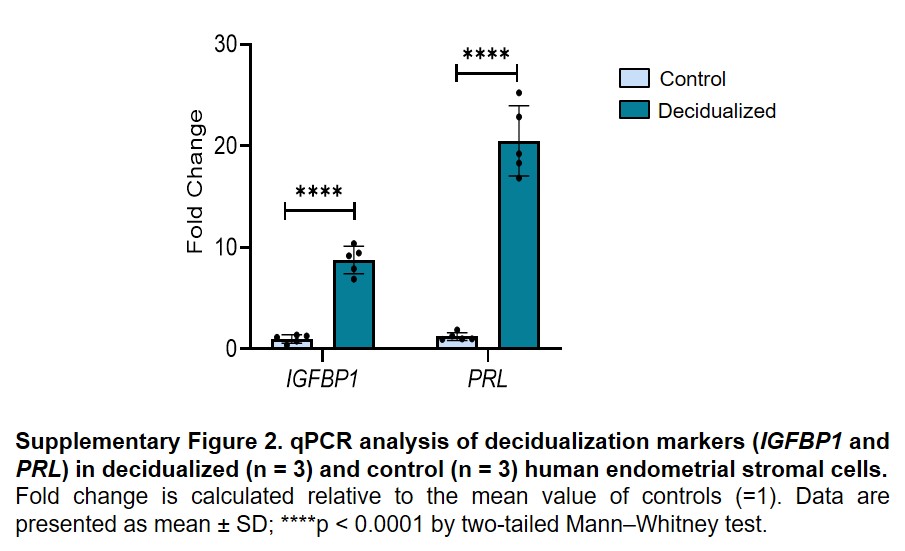

### Supplymental Fig 3

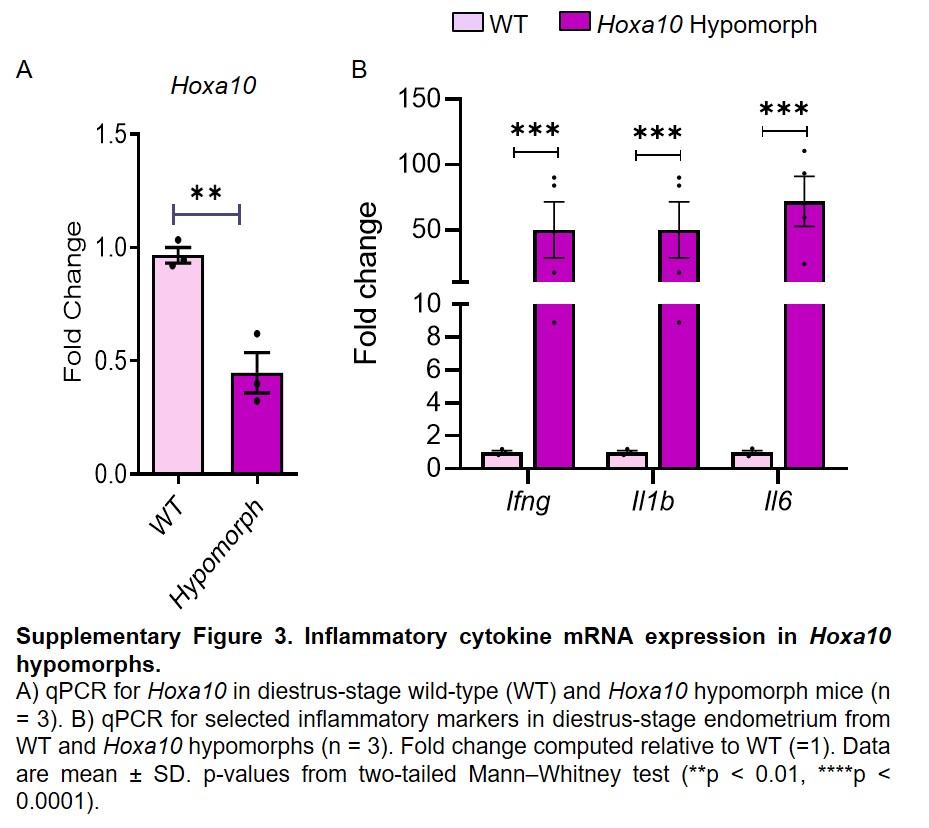

### Supplymental Fig 4

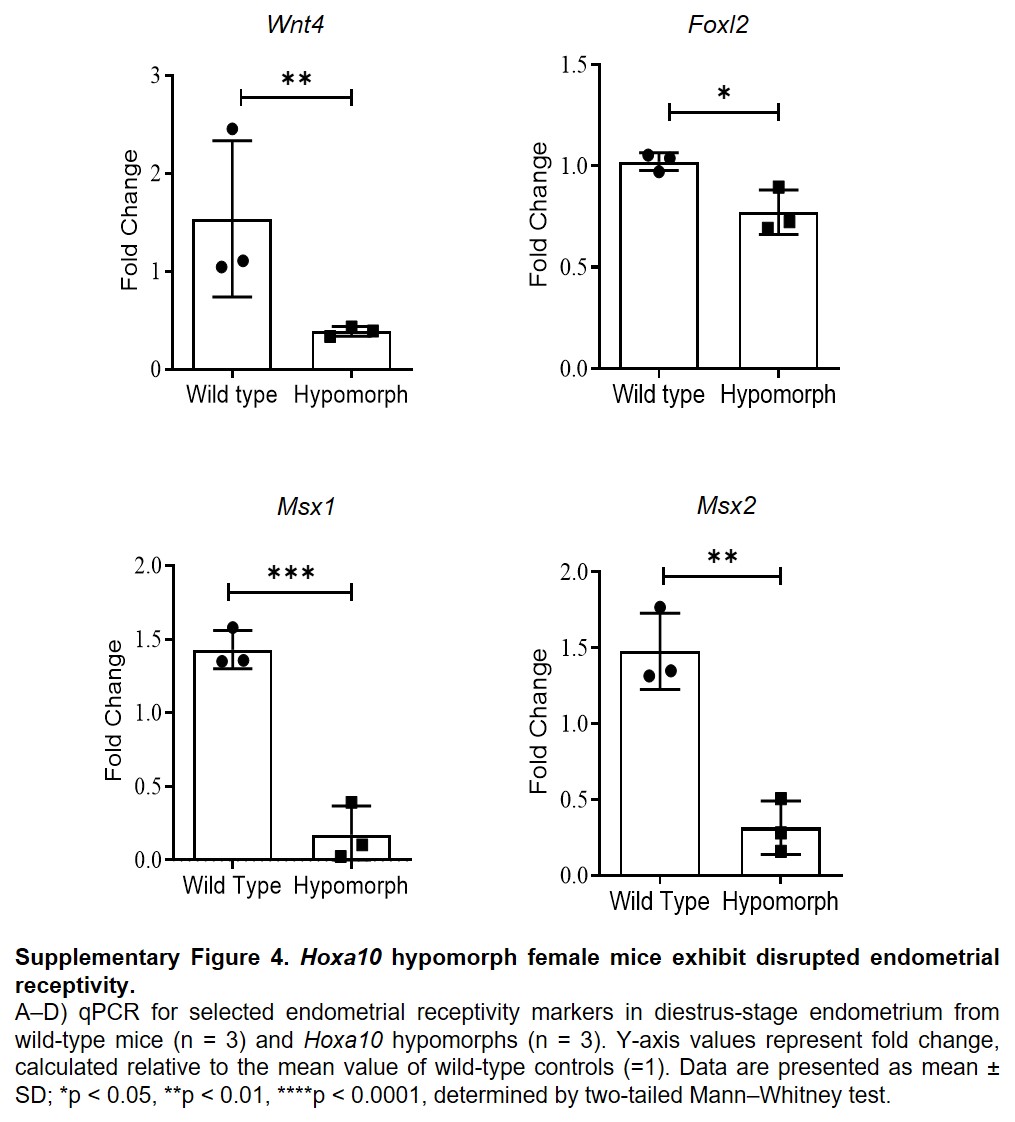

### Supplymental Fig 5

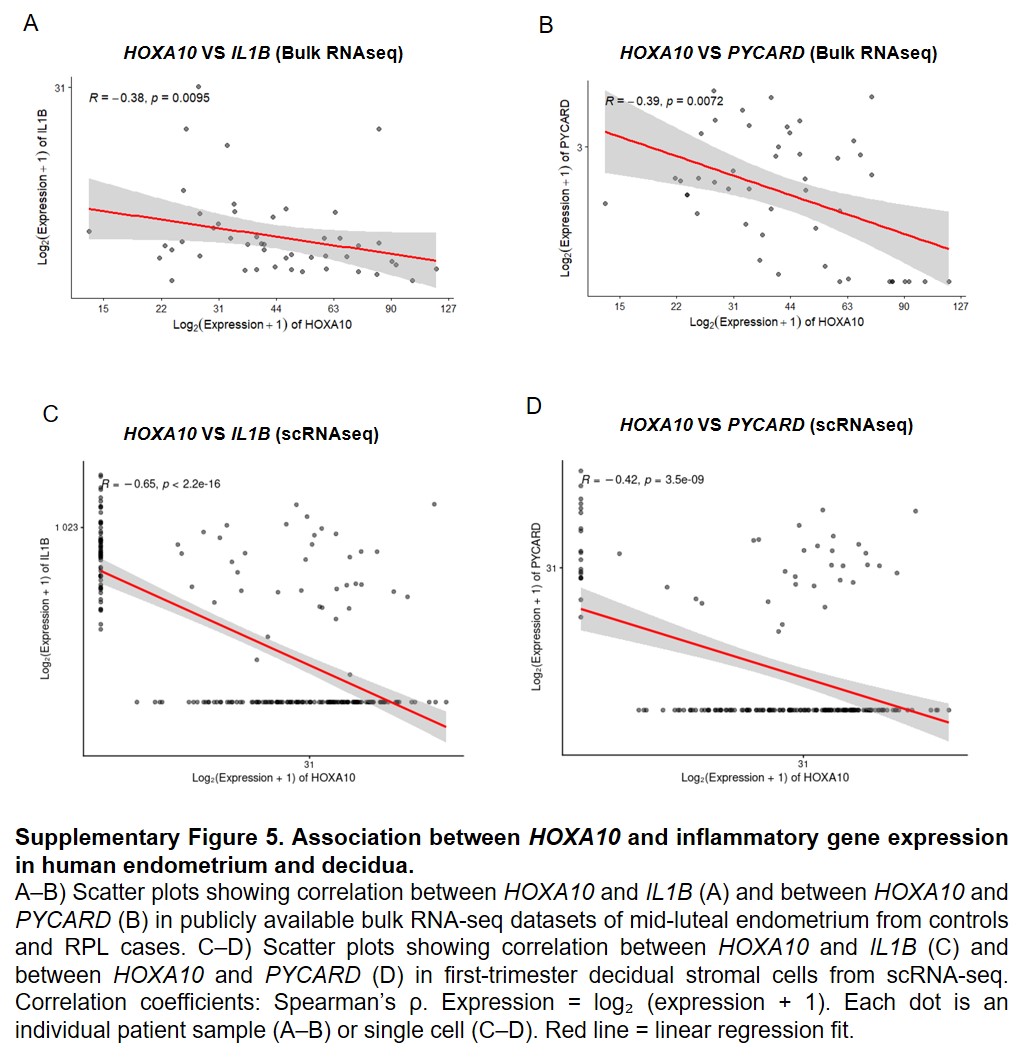
