## Supplementary material for "Temporal Control of Decidual Inflammation by HOXA10 is Essential for Implantation and its Dysregulation is Associated with Early Pregnancy Loss": Supplymental table 1-3

Supplementary Table 1: RT-qPCR Primers used in the present study

| **Gene Name** | **Primer sequence (5’-3’)** | **Annealing Temperature (°C)** | **Product Size (bp)** |
| --- | --- | --- | --- |
| *Foxl2* | F 5’ GGCTCTTCGGGAGCGGAGGA  3’  R 5’ TGGCAGGAGGCGTAGGGCAT 3’ | 61 | 163 |
| *GSN* | F 5’ CTTTGTCTGGAAAGGCAAGC 3’  R 5’ GCGATATGGCTGGAAAGGTA 3’ | 60 | 219 |
| *Hoxa10* | F 5’ ACTAAGAGCAGCACGGTACG 3’  R 5’ CCTTTGGAACTGCCTTGACTC 3’ | 58 | 200 |
| *HOXA10* | F 5’ GCCCCTTCCGAGAGCAGAAAA G 3’  R 5’AGGTGGAGCCTGCGGCTAATCTCTA 3’ | 65 | 211 |
| *IGFBP1* | F 5’ GAGTTTAGCCAAGGCACAGG 3’  R 5’ GAGACCCAGGGATCCTCTTC 3’ | 60 | 173 |
| *Ifng* | F 5’ GGCCATCAGCAACAACATAAGCGT 3’  R 5’ TGGGTTGTTGACCTCAAACTTGGC 3’ | 118 | 62 |
| *IL11* | F 5’ CTGAGCCTGTGGCCAGATA 3’  R 5’ AGCTGTAGAGCTCCCAGTGC 3’ | 66 | 236 |
| *IL13* | F 5’ GTACTGTGCAGCCCTGGAAT 3’  R 5’ TTTACAAACTGGGCCACCTC 3’ | 65 | 162 |
| *IL15* | F 5’ AGAAGCCAACTGGGTGAATG 3’  R 5’ TACTTGCATCTCCGGACTCA 3’ | 62 | 191 |
| *IL17A* | F 5’ ACCAATCCCAAAAGGTCCTC 3’  R 5’ GGGGACAGAGTTCATGTGGT 3’ | 60 | 171 |
| *IL1B* | F 5’ GGGCCTCAAGGAAAAGAATC 3’  R 5’ TTCTGCTTGAGAGGTGCTGA 3’ | 57 | 204 |
| *IL6* | F 5’ AGGAGACTTGCCTGGTGAAA 3’  R 5’ CAGGGGTGGTTATTGCATCT 3’ | 59 | 180 |
| *ITGA5* | F: 5’ CGAGACCTGGATGGCAATG 3’  R: 5’ TCTGGTTCACGGCAAAGTAGTC 3’ | 53 | 786 |
| *ITGA6* | F: 5’ TTGGGCGGTGTTATGTTCTG 3’  R: 5’ CACCCATCCTTGTTGAGGTCC 3’ | 52 | 527 |
| *ITGAV* | F: 5’ ACTGGGAGCACAAGGAGAACC 3’  R: 5’ CCGCTTAGTGATGAGATGGTC 3’ | 53 | 290 |
| *LIF* | F 5’ CTGTTGGTTCTGCACTGGAA 3’  R 5’ GCCACATAGCTTGTCCAGGT 3’ | 59 | 216 |
| *Msx1* | F 5’ TCTCGGCCATTTCTCAGTCG 3’  R 5’ GGGACTCAGCCGTCTGGC 3’ | 60 | 145 |
| *Msx2* | F 5’ CTCTCGTCAAGCCCTTCGAG 3’  R 5’ CTCATATGTCTGGGCGGCG 3’ | 60 | 109 |
| *PRL* | F 5’ TGCAGATGGCTGATGAAGAG 3’  R 5’ TGCAATGGAACGGATCATTA 3’ | 62 | 175 |
| *Wnt4* | F 5’ TGGACTCCCTCCCTGTCTTTGGGA 3’  R 5’ TCCTGACCACTGGAAGCCCTGTG 3’ | 64 | 188 |
| *18S* | F 5’ GGAGAGGGAGCCTGAGAAAC 3’  R 5’ CCTCCAATGGATCCTCGTTA 3 | 60 | 180 |
| *HOXA10*  siRNA | Duplex1: CTCGTCCTCTTTCGCGCAGAA  Duplex2: TACTGTGAAGTTACATGCATA | - | - |
| Scrambled siRNA | Qiagen, catalog no. 1027101 |  |  |

Supplementary Table 2: List of antibodies and their optimized dilutions used in this study

| **Primary Antibody** | **Source** | **Catalogue No.** | **Dilution** |
| --- | --- | --- | --- |
| ASC | Abclonal | A11433 | 1:100 |
| Caspase1 | Abclonal | A0964 | 1:100 |
| HOXA10 | In-house | * | 1:25 |
| IL1β | Abclonal | A11370 | 1:100 |
| NLRP3 | Abclonal | A5652 | 1:100 |
| TXNIP | Abclonal | A9342 | 1:100 |

Note:

*: In-house HOXA10 antibody was validated and the results can be found at, doi: <https://doi.org/10.1101/2025.01.10.631632> (Ashary et al., 2025)

Supplementary Table 3: Summary of human transcriptomic datasets used in this study

| Accession ID | Type of data | Study group | Tissue type | Stage |
| --- | --- | --- | --- | --- |
| GSE65099 / PRJNA273001 | Bulk RNA-seq | Control and RPL | Endometrial Biopsy | Mid-Luteal Phase (LH+6 - 10) |
| PRJNA314429 | Bulk RNA-seq | Control and RPL | Endometrial Biopsy | Mid-Luteal Phase (LH+6 - 10) |
| GSE86491 / PRJNA342633 | Bulk  RNA-seq | Control | Endometrial Biopsy | Mid-Luteal Phase (LH+6 - 10) |
| GSE98386 / PRJNA384963 | Bulk RNA-seq | Control | Endometrial Biopsy | Mid-Luteal Phase (LH+6 - 10) |
| PRJEB25794 | scRNA-seq | Control | Decidua | First trimester decidua (6 to 12 weeks) |
| OEP002901 | scRNA-seq | Control and RPL | Decidua | First trimester decidua (7 to 9 weeks) |
